# Targeting ATP12A, a non-gastric proton pump alpha subunit, for idiopathic pulmonary fibrosis treatment

**DOI:** 10.1101/2022.06.08.495330

**Authors:** Mohamed Abdelgied, Katie Uhl, Oliver G Chen, Chad Schultz, Kaylie Tripp, Angela M Peraino, Shreya Paithankar, Bin Chen, Maximiliano Tamae Kakazu, Alicia Castillo Bahena, Tara E Jager, Cameron Lawson, Dave W Chesla, Nikolay Pestov, Nikolai N. Modyanov, Jeremy Prokop, Richard R Neubig, Bruce D Uhal, Reda E Girgis, Xiaopeng Li

## Abstract

Idiopathic Pulmonary Fibrosis (IPF) is a pathological condition of unknown etiology which results from injury to the lung and an ensuing fibrotic response that leads to thickening of the alveolar walls and obliteration of the alveolar space. The pathogenesis is not clear and there are currently no effective therapies for IPF. Small airway disease and mucus accumulation are prominent features in IPF lungs, similar to Cystic Fibrosis (CF) lung disease. The ATP12A gene encodes the alpha-subunit of the non-gastric H^+^, K^+^-ATPase, which functions to acidify the airway surface fluid and impairs mucociliary transport function in cystic fibrosis patients. We hypothesize that the ATP12A protein may play a role in the pathogenesis of IPF. Our studies demonstrate that ATP12A protein is overexpressed in distal small airways from IPF patient lungs compared to normal human lungs. In addition, overexpression of the ATP12A protein in mouse lungs worsened the bleomycin (BLEO)-induced experimental pulmonary fibrosis. This was prevented by a potassium-competitive proton pump blocker, vonoprazan (VON). This data supports the concept that the ATP12A protein plays an important role in the pathogenesis of lung fibrosis. Inhibition of the ATP12A protein has the potential as a novel therapeutic strategy in IPF.

## Introduction

Approximately 50,000 new patients are diagnosed with idiopathic pulmonary fibrosis (IPF) annually, and approximately 40,000 patients die every year from IPF in the USA (1). IPF is a serious health condition that impairs the ability of the lung to exchange oxygen. The exact cause of IPF is unknown, but an injury to the lung can lead to the buildup of fibrotic tissue within the air sacs; as the disease progresses, structures that are crucial to the absorption of oxygen are eventually destroyed (2, 3). The FDA-approved drugs Pirfenidone and Nintedanib can slow the decline in lung function but cannot reverse the damage that has already been done to the tissue; morbidity and mortality remain high (4, 5). Hence, novel therapeutic targets are urgently needed.

The mechanisms underlying the pathogenesis of IPF are unclear (6–8). The main pathologic features include the collapse and obliteration of distal small airways, the proliferation and accumulation of fibroblast/myofibroblast cells, and increased collagen deposition in the alveolar spaces (9, 10). Recent evidence suggests that distal small airways are involved in the early pathogenesis of IPF. Epithelial microfoci of injury and a failure of reepithelization caused by the interplay of genetic predisposition, aging, and environmental factors contribute to fibroblast activation and infiltration of the alveolar spaces (9, 11–13). Small airways are defined as having a diameter less than 2 mm in adult human lungs (14). There is active bronchiolar remodeling, including collagen deposition, found in small airways of IPF patients (15–17). Studies on lung explants from IPF patients compared to controls demonstrate that approximately 57% of terminal bronchioles are lost in regions of minimal fibrosis, which is associated with the appearance of fibroblastic foci (18, 19). This suggests that loss of small airways may precede fibrotic changes (18, 19). In addition, Mucin 5B (MUC5B), the gene for a major gel-forming mucin, accumulates in distal small airways in IPF patients. Overexpression of MUC5B leads to mucus accumulation in small airways and enhances bleomycin (BLEO)-induced lung fibrosis (20–22). However, the mechanism behind mucus accumulation in IPF distal lung airways is unclear.

Mucus accumulation is also the main manifestation of cystic fibrosis (CF) lung disease caused by mutations in the cystic fibrosis transmembrane conductance regulator (CFTR) gene. There is a mucociliary transport (MCT) defect in CF airways due to lower airway surface liquid (ASL) pH. In the current study, we first investigated ATP12A protein expression levels in IPF, COPD, and normal human airways. ATP12A genes (previously termed ATP1AL1) encode the catalytic α-subunits of the non-gastric H-K-ATPases (ngHKA)(23). Non-gastric H-K-ATPase (ngHKA) represent a third group of potassium-dependent ion-transporting P-type ATPases that are equally distinct from closely related Na, K-ATPases, and gastric H, K-ATPases in structure-functional properties (24–26). ATP12A (the catalytic α-subunit) assembles with β-subunit (ATP1B1) forming an active ion pump (ngHKA) in apical membranes of epithelial cells of many different tissues including colon, kidney, skin, penis, lung, pancreas, etc (25, 27–30). The ATP12A protein functions as the alpha subunit of a proton pump that acidifies ASL (31). For this study, the use of the term “ATP12A” refers to the α-subunit. We hypothesize that ATP12A-mediated acid secretion in IPF distal lung small airways may regulate ASL pH, leading to lower ASL pH and impaired MCT, thus contributing to IPF pathophysiology. To test this hypothesis, we evaluated ATP12A gene expression in IPF lungs and tested its function in primary cell cultures and animal models. Our findings in this study elucidated a potentially important yet previously underappreciated role of small airway epithelia in the pathogenesis of IPF. Understanding the contribution of small airways and ASL pH to IPF pathogenesis may lead to the discovery of a new therapeutic strategy.

## Results

### Increased ATP12A expression in the small airways of IPF patient lungs

ATP12A is demonstrated to acidify the airway lining fluid, impair mucociliary transport function in CF patients, and is upregulated in CF large airways (32). We previously reported that ATP12A is not expressed in small airways from normal human distal lungs (33). To test if ATP12A is expressed in IPF distal lungs, we investigated the expression of ATP12A in lung explant samples collected from advanced IPF (N=13) and chronic obstructive pulmonary disease (COPD, N=8) patients undergoing lung transplantations. Before enrollment into this study, pathological analysis was carried out to confirm a diagnosis of usual interstitial pneumonia (UIP) in the IPF samples and emphysema in the COPD samples. Normal lung control samples (N=4) were collected from donor lungs that did not meet the criteria for transplantation. After dissection of the lungs, samples were fixed and processed for detection of ATP12A, MUC5B, and Mucin 5AC (MUC5AC) by immunofluorescence, immunohistochemistry, and in situ hybridization. ATP12A overexpression was detected in the large airways and submucosal glands (SMG) of both IPF and COPD, and the small airways of IPF lungs. Normal lung large airways, submucosal glands, and small airways showed a low level of ATP12A expression (**Figure 1A and B; Supplementary Figure 1A and B**). The mean relative staining intensity of ATP12A was 15.6± 3.6 in the normal large airway, 78.4 ± 6.4 in IPF large airways, and 70.5± 9.6 in COPD large airways. The mean relative staining intensity of ATP12A was 12.9 ± 4.1 in normal lung SMG, 59.5+6.8 in IPF lung SMG, and 41.9± 7.4 in COPD lung SMG. The mean relative staining intensity of ATP12A was 5.2± 3.0 in the normal small airway, 66.8± 6.4 in IPF small airways, and 9.4± 2.3 in COPD small airways. Also, both IPF and COPD patients’ large airways, submucosal glands, and small airways showed an increased expression of MUC5B and MUC5AC compared to those of the normal lungs (**Figure 1C, Supplementary Figure 1C**). The surface epithelium of the IPF lungs’ small airways showed overexpression of ATP12A associated with MUC5B accumulation (**Figure 1C**). Those data demonstrated that ATP12A is overexpressed and colocalized with Muc5B accumulation in IPF distal lungs.

**Figure 1.**
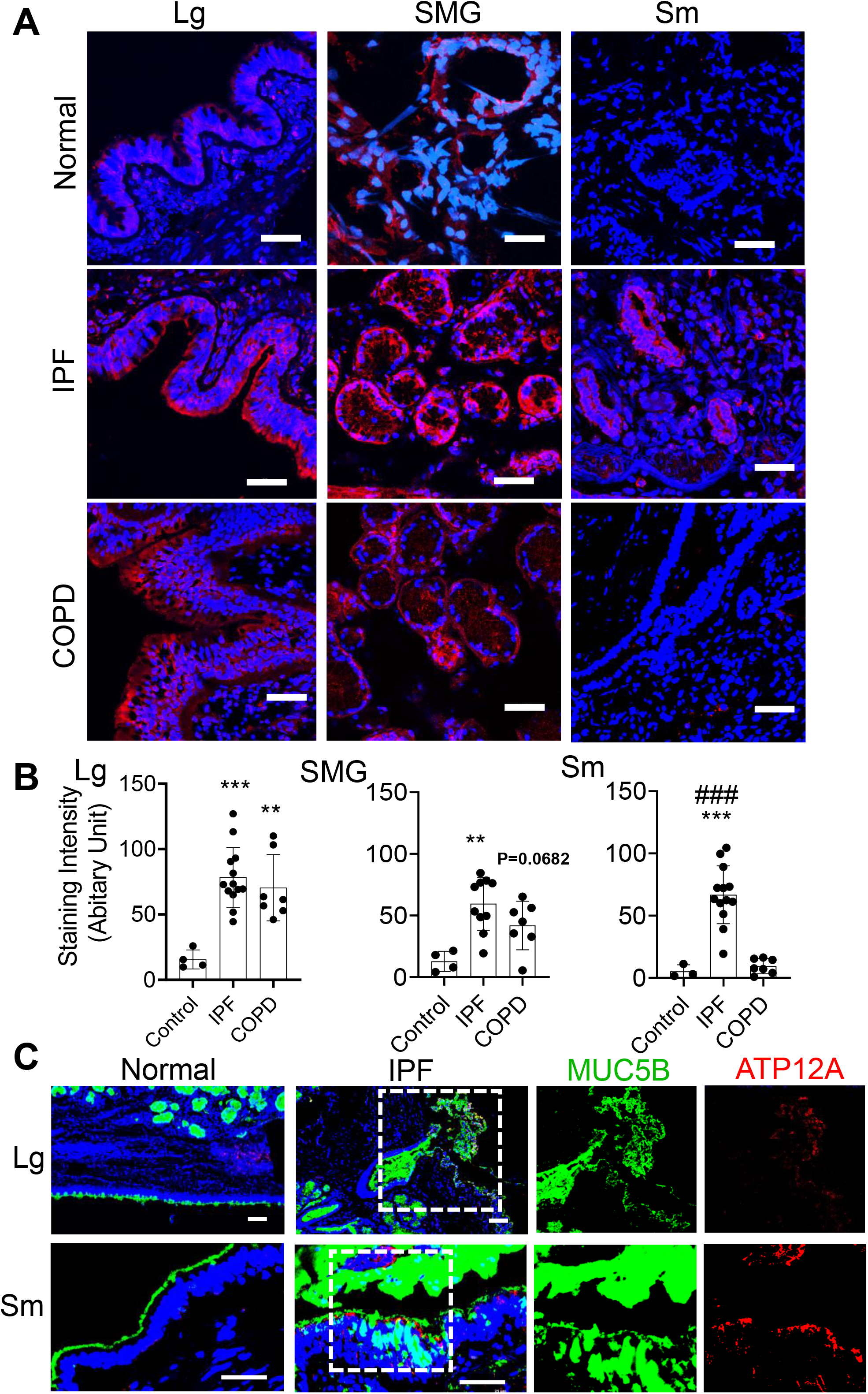
ATP12A and MUC5B proteins expression in human lung explants. **(A)** Representative confocal microscope images showing immunodetection of ATP12A (red) by immunofluorescence. Nuclei were counterstained by DAPI (blue). Scale bar: 25 µm. Images show large airways (Lg), submucosal glands (SMG), and small airways (SM) of normal (upper panel), IPF (middle panel), and COPD (lower panel) human lungs. ATP12A overexpression was found in large airways and submucosal glands of IPF and COPD, and in the small airways of IPF lungs. **(B)** ATP12A immunofluorescence staining intensity quantification charts. Data expressed as mean ± STDEV of 4 normal, 13 IPF, and 8 COPD human lung samples. **,*** indicates p-value < 0.01 and 0.001 compared to control, respectively. ### indicates p-value < 0.001 compared to COPD. **(C)** Representative confocal microscope images showing immunodetection of ATP12A (red) and MUC5B (green) in large airways (Lg) and small airways (SM) of normal and IPF human lungs. Nuclei were counterstained by DAPI (blue). Scale bar: 25 µm. ATP12A and MUC5B were overexpressed in both large and small airways of IPF lungs.

### Induced expression of ATP12A in mice airways worsened bleomycin-induced pulmonary fibrosis

Murine airways lack ATP12A expression (31). To mimic the ATP12A expression found in humans, we induced ATP12A expression in mouse airways via intratracheal instillation of adenovirus subtype 5 encoding mouse ATP12A (Ad-ATP12A). Adenovirus subtype 5 encoding GFP (Ad-GFP) was administrated intratracheally in control mice. 15 days after instillation, ATP12A expression in mice airways was confirmed by RT-PCR (**Figure 2B**), in situ hybridization (**Figure 2A 2 and 4**), and immunofluorescence staining (**Figure 2A 1 and 3**). ATP12A expression was detected on the apical surface of mouse airway epithelium as well as in the lung parenchyma (**Figure 2A 3**). Additionally, ATP12A mRNA molecules were found in the airway epithelium (**Figure 2A 4**). In contrast, no ATP12A was detected in Ad-GFP-infected mouse lungs, but GFP protein was expressed in mouse airways (**Supplementary Figure 2**).

**Figure 2:**
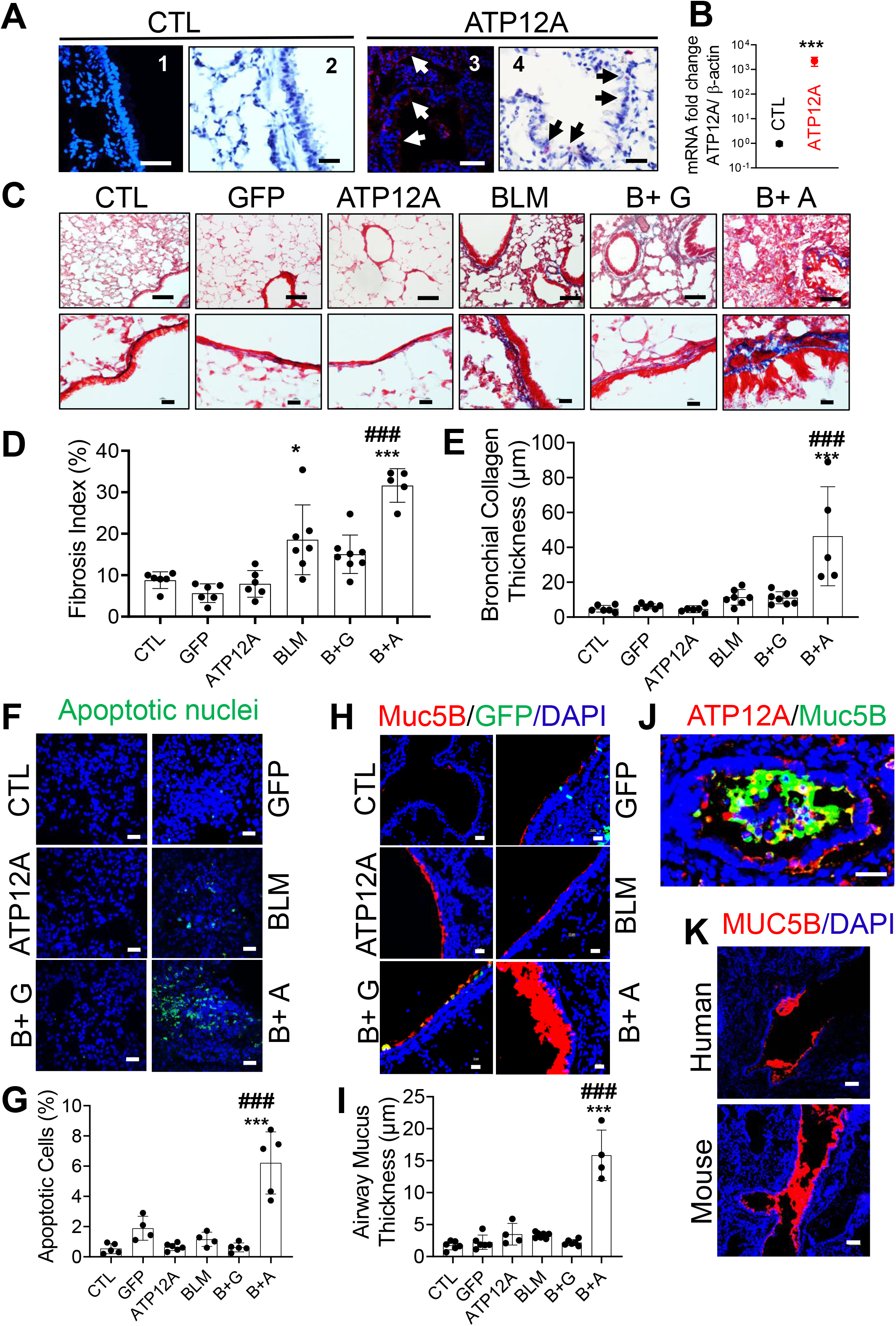
Induced expression of ATP12A in mice airways worsens bleomycin-induced pulmonary fibrosis after intratracheal exposure. **(A)** Confocal and brightfield microscopes images of control and Ad-ATP12A treated mice lungs showing expression of ATP12A in mice airways; **(A 1, 3)**. Confocal microscope images show immunodetection of ATP12A (red, white arrows) by immunofluorescence. Nuclei were counterstained by DAPI (blue). Scale bar: 25 µm. **(A 2, 4)**. brightfield microscope images ATP12A mRNA detection (pink, black arrows) by in situ hybridization (ISH). Nuclei were counterstained by hematoxylin (light blue). Scale bar: 25 µm. **(B)** Quantitative RT-PCR analysis of ATP12A gene expression level in mice lungs. Data are expressed as mean ± STDEV. **(C)** Bright field microscope images of mice lung tissue stained with Masson’s trichrome showing collagen deposition (blue) in the lung, with the upper panel showing collagen deposition throughout the lung. Scale bar 100 µm; lower panel shows collagen deposition in the peribronchial region. Scale bar 10 µm. **(D)** Chart shows the fibrosis index in the experimental mouse lungs. Data are expressed as mean ± STDEV. n≥5 animals/group. **(E)** Chart shows parabronchial collagen thickness (µm) in mouse lungs. Data are expressed as mean ± STDEV. n≥5 animals/group. **(F)** Confocal microscope images show cellular apoptosis of lung epithelial cells by TUNEL staining. Apoptotic cell nuclei are stained green. Scale bar: 25 µm. **(G)** Chart shows the percentage of apoptotic cells in mouse lungs. Data are expressed as mean ± STDEV, with n≥5 animals/group. **(H)** Confocal microscope images show immunodetection of MUC5B (red) by immunofluorescence. Nuclei were counterstained by DAPI (blue). Scale bar: 25 µm. **(I)** Chart shows airway mucus thickness in mouse lungs. Data are expressed as mean ± STDEV. n≥5 animals/group. **(J)** Confocal microscope image show immunodetection of ATP12A (red) and MUC5B (green) by immunofluorescence in mice lung airway. Nuclei were counterstained by DAPI (blue). Scale bar: 25 µm. **(K)** Confocal microscope images show immunodetection and accumulation of MUC5B (red) in human (upper panel) and mice (lower panel) airways by immunofluorescence. Nuclei were counterstained by DAPI (blue). Scale bar: 25 µm. CTL: untreated control group; GFP: Adenovirus expressing green fluorescence (Ad-GFP) treated group; ATP12A: Adenovirus expressing mouse ATP12A (Ad-ATP12A) treated group; BLM: Bleomycin treated group; B+G: Bleomycin and Ad-GFP treated group; B+A: Bleomycin and Ad-ATP12A treated group. *,*** indicates p-value < 0.05 and 0.001 compared to control, respectively. ### indicates p-value < 0.001 compared to bleomycin treated group.

To determine whether ATP12A plays a role in pulmonary fibrosis, we injured mice with BLEO and assessed the intensity of pulmonary fibrosis indicators. Mice expressing ATP12A showed a significant increase in collagen deposition in the lung (**Figure 2C, D, and E**), apoptosis in the alveolar epithelium, (**Figure 2F and G; Supplementary Figure 2**), and accumulation of mucus in airways (**Figure 2H, I, and J**) compared to mice exposed only to BLEO. Mice expressing ATP12A in the lung showed extensive collagen deposition in the lung parenchyma, especially in areas adjacent to the bronchi, with a significant increase in collagen deposition in the peribronchial area (**Figure 2C**). Mucus accumulation and airway blockage were also observed in mice expressing ATP12A and challenged with BLEO, which mimicked the mucus accumulation observed in human IPF lung airways (**Figure 2K**). ATP12A expression in the apical surface of mouse airways epithelium was associated with airway mucus accumulation, **(Figure 2J**), suggesting that ATP12A has a role in this process. Interestingly, induced ATP12A expression in mouse lungs stimulated early honeycombing cyst formation after BLEO exposure (**Figure 5H and I**).

**Figure 3:**
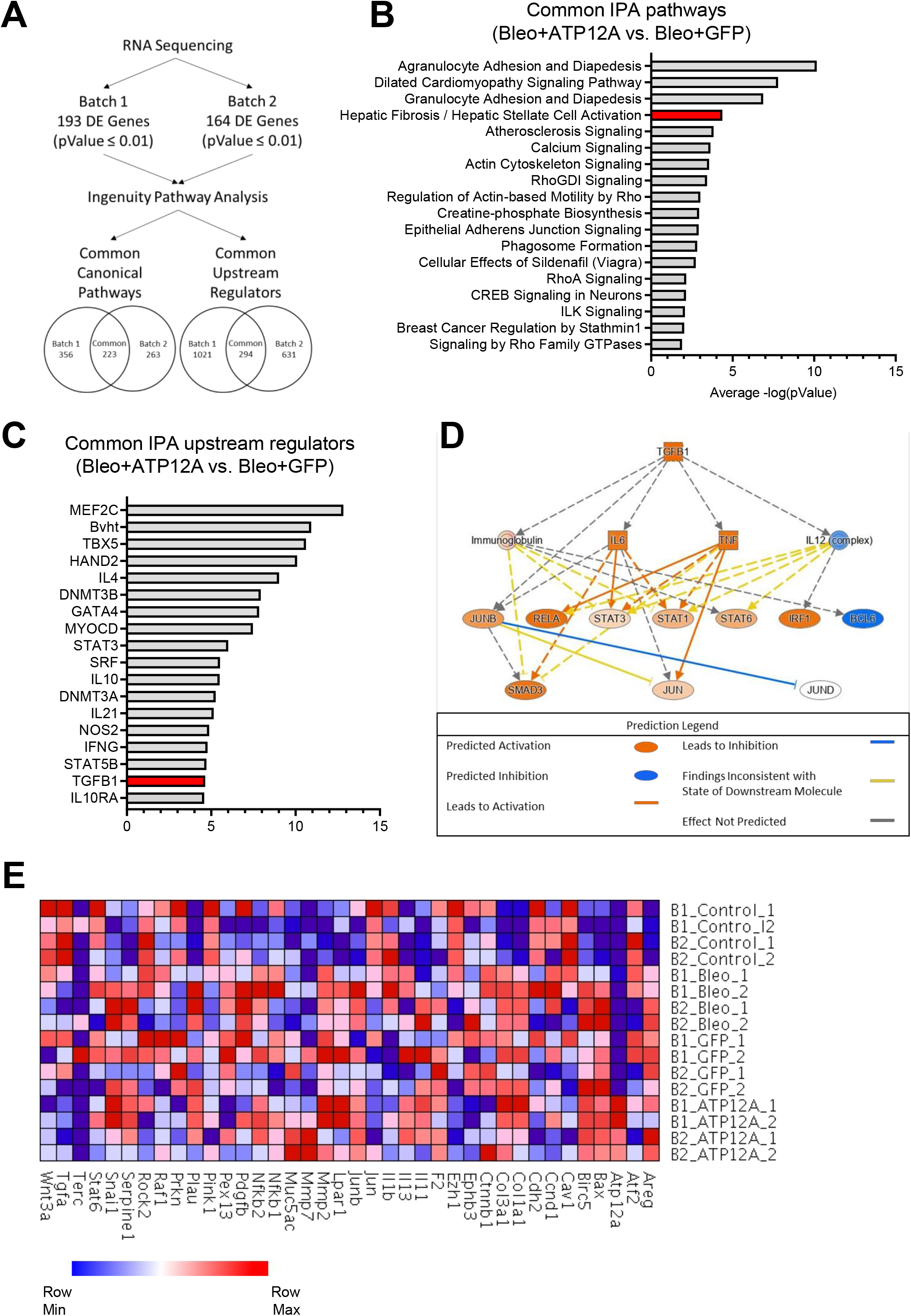
Overexpression of ATP12A in mouse lungs enhanced fibrotic pathway and TGF-β1 signaling pathway in bleomycin-induced pulmonary fibrosis. **(A)** Diagram showing the data analysis workflow. Data collected from the bulk RNA sequencing of mouse lung tissue following treatment with bleomycin and the introduction of ATP12A or GFP via adenovirus delivery. Sequencing and differential expression analysis were performed on two batches of samples (pValue = 0.01). The lists of DE genes were submitted to the QIAGEN Ingenuity Pathway Analysis (IPA) to identify canonical pathways and upstream regulators. **(B)** The common canonical pathways from the Bleo/ATP12A vs. Bleo/GFP comparison in each batch were compiled and arranged according to the average -log(pValue), as calculated by IPA. The hepatic fibrosis/hepatic stellate cell activation pathway has been highlighted in red. **(C)** Common upstream regulators were identified between the Bleo/ATP12A vs. Bleo/GFP comparison in the two batches and arranged based on the average -log(pValue) calculated by IPA. The TGF-β1 upstream regulator has been highlighted in red. **(D)** Pathway diagram displaying the predicted activation states of molecules in the TGF-β1 signaling pathway based on differential gene expression data submitted to IPA. **(E)** Heatmap displaying the expression levels of selected genes from the idiopathic pulmonary fibrosis pathway, as listed by IPA, as well as genes of interest included by the authors (Atp12A and Muc5ac).

**Figure 4.**
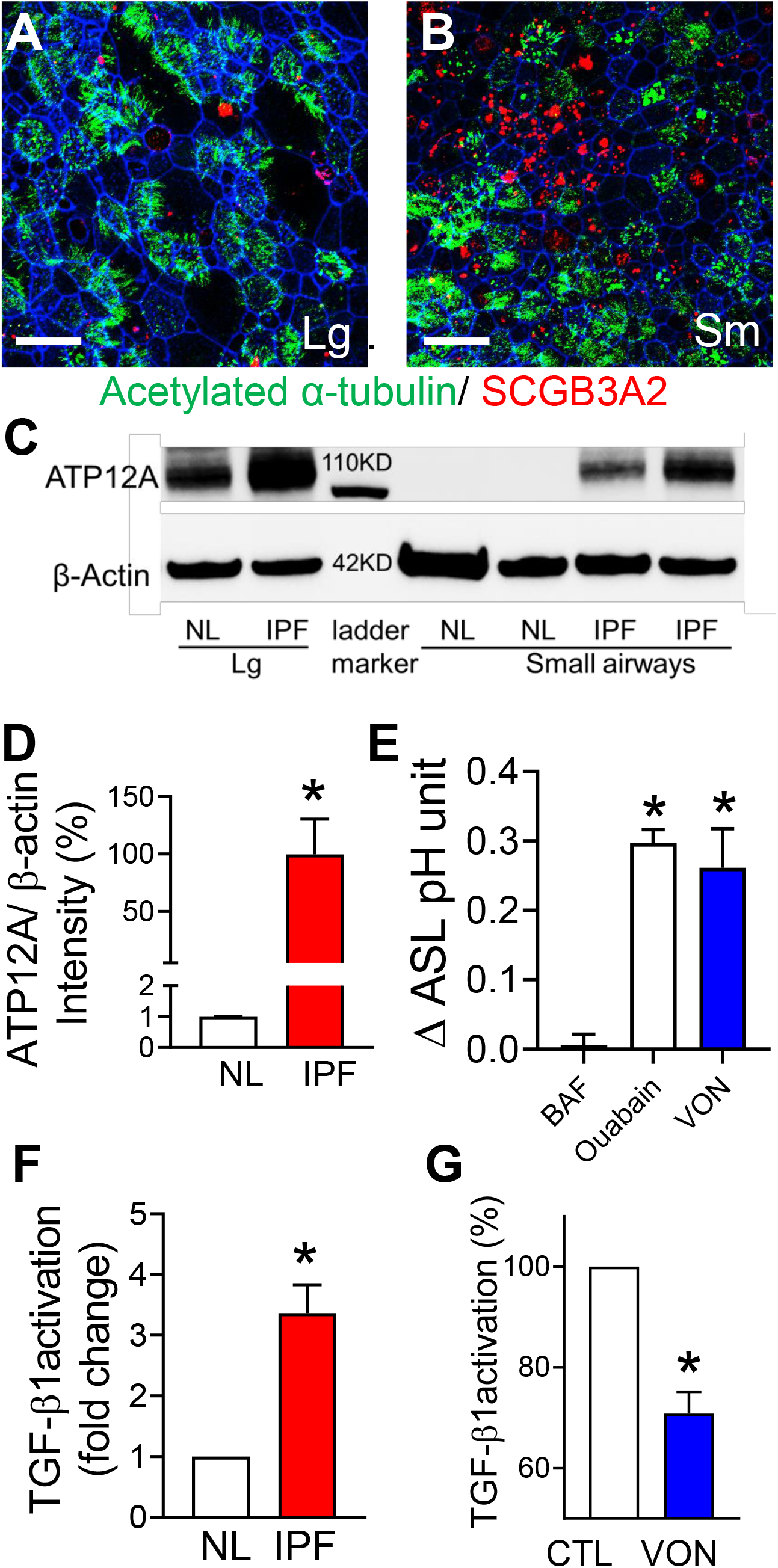
ATP12A expression was increased in IPF small airways in vitro. Human large. **(A)** and small **(B)** airway epithelial culture. Scale bar: 25 µM. SCGB3A2 in red, acetylated α-tubulin in green, F-actin in blue. **(C)** ATP12 expression was detected by immunoblotting in culture large and small airways cells from normal (NL) and IPF lungs. **(D)** Semi-quantification of band intensity showed that ATP12A expression in IPF small airways culture was increased about 100 folds, compared to normal lungs (NL). N=8, * P<0.05. **(E)** Ouabain and VON increased ASL pH; but Bafilomycin (BAF) has no effect on ASL pH in IPF small airway. N=5, * P<0.05. **(F)** IPF small airway apical surface activated more latent TGF-β1. N=3, * P<0.05. **(G)** VON decreased 30% of TGF-β1 activation in IPF small airways. N=6, * P<0.05.

**Figure 5:**
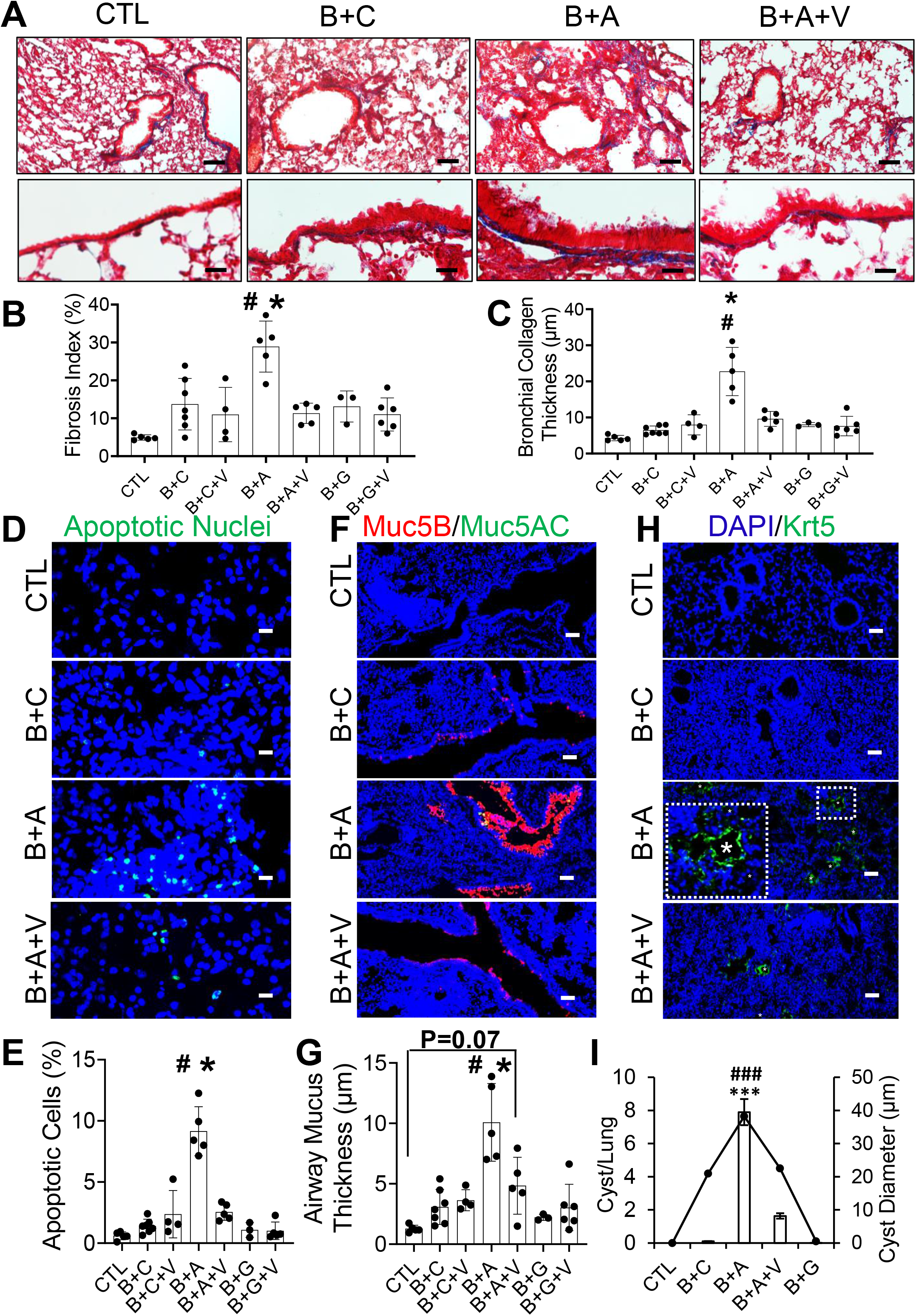
Inhibition of ATP12A by potassium competitive proton pump blocker, VON, reduced bleomycin-induced pulmonary fibrosis in mice expressing ATP12A. **(A)** Bright field microscope images of mice lung tissue stained with Masson’s trichrome showing collagen deposition (blue) in the lung; upper panel shows collagen deposition throughout the lung. Scale bar 100 µm; lower panel shows collagen deposition in the peribronchial area. Scale bar 10 µm. **(B)** Chart shows fibrosis index in mice lung. Data are expressed as mean ± STDEV, with n≥5 animals/group. **(C)** Chart shows parabronchial collagen thickness (µm) in mice lungs. Data are expressed as mean ± STDEV; n≥5 animals/group. **(D)** Confocal microscope images show cellular apoptosis of lung epithelial cells by TUNEL staining. Apoptotic cell nuclei are stained green. Scale bar: 25 µm. **(E)** Chart shows the percentage of apoptotic cells in mouse lungs. Data are expressed as mean ± STDEV; n≥5 animals/group. **(F)** Confocal microscope images show immunodetection of MUC5B (red) and MUC5AC (green) by immunofluorescence. Nuclei were counterstained by DAPI (blue). Scale bar: 25 µm. **(G)** Chart shows airways mucus thickness in mouse lung. Data are expressed as mean ± STDEV; n≥5 animals/group. **(H)** Confocal microscope images show immunodetection of keratin 5(Krt5, basal cell marker in green) by immunofluorescence. Inset shows a honeycomb cyst (white asterisk) with Krt5+ cell lining (green). Nuclei were counterstained by DAPI (blue). Scale bar: 25 µm. **(I)** Chart shows the number of honeycomb cysts per right middle lung lobe (column chart) and the average diameter of honeycomb cyst (linear plot). Data are expressed as mean ± STDEV; n≥3 animals/group. CTL: untreated control group; B+C: Bleomycin and carboxymethyl cellulose treated group; B+C+V: Bleomycin, carboxymethyl cellulose, and VON fumarate treated group; B+A: Bleomycin and Ad-ATP12A treated group. B+A+V: Bleomycin, Ad-ATP12A, and VON fumarate treated group. B+G: Bleomycin and Ad-GFP treated group; B+G+V: Bleomycin, Ad-GFP, and VON fumarate treated group. *** indicates p-value < 0.001 compared to control; ### indicates p-value < 0.001 compared to B+A+V group.

### Overexpression of ATP12A in mouse lungs enhanced fibrotic pathway and TGF-β1 signaling pathway in bleomycin-induced pulmonary fibrosis

To investigate the mechanism of how ATP12A enhances pulmonary fibrosis, bulk RNAseq was performed to evaluate the mRNA expression in BLEO-induced lung fibrosis in mouse lungs treated with Ad-ATP12A or Ad-GFP. The bulk RNA-sequencing data for the mouse lung tissue samples were divided into two groups to reduce the batch effect (**Figure 3A**). Each group consisted of eight mice, with two mice per treatment group. Differential expression analysis of Batch 1 revealed 193 DE genes with a p-value ≤ 0.01; Batch 2 had 164 DE genes with the same p-value threshold. The gene list for the individual batches was submitted to Ingenuity Pathway Analysis to identify common canonical pathways and common upstream regulators. Batch 1 returned a total of 356 canonical pathways and the data from Batch 2 represented 263 pathways; there were 223 canonical pathways shared between the batches. The same comparison was performed for the upstream regulators, with 294 common regulators identified.

Mice lungs that had the adenovirus-induced expression of ATP12A and were challenged with BLEO demonstrated an increase in pathways that are known to play a role in the development of fibrosis. According to the IPA canonical pathway analysis, at least ten genes are shared between the hepatic fibrosis/hepatic stellate cell activation pathway and the pulmonary fibrosis pathway (**Figures 3B and E**). The shared genes include ACTA2, a key regulator of myofibroblast development, COL1A1, and the transcription factor JUN. These genes are all found to have elevated expression levels in pulmonary fibrosis. Also related to the fibrosis pathway is the actin cytoskeleton signaling pathway, which plays an important role in myofibroblast differentiation. The ILK signaling pathway is involved in the up-regulation of the filament protein vimentin, the inhibition of which protects the lungs from fibrotic injury (34, 35). One of the most noteworthy upstream regulators shared by the two batches of samples is TGF-β1. This protein plays a role in the production of the extracellular matrix, and increased levels of TGF-β1 expression have been reported in pulmonary fibrosis (**Figures 3C and D**). Interleukin 4 (IL4) was also identified as a common upstream regulator, as well as STAT3. Like TGF-β1, expression levels of IL4 and STAT3 are increased in pulmonary fibrosis patients (36, 37). In addition, analysis of the gene expression data for Ad-ATP12A treated samples revealed the activation of the TGF-β1 signaling pathway. The IPA analysis tool uses its extensive literature curated database to determine key regulator molecules in a signaling pathway. The TGF-β1 signaling pathway was overlaid with the gene expression data from the two sample batches to predict the activation states of the proteins involved in the pathway (**Figure 3D**). The IL12 complex and BCL6 are the only proteins predicted to be inhibited in the BLEO/ATP12A vs. BLEO/GFP dataset. TGF-β1, IL6, TNF, as well as several members of the STAT family are predicted to be activated. There was not enough power for the analysis tool to predict the state of JUND; however, JUNB, a known inhibitor of JUND, is predicted to be activated. Overall, the treatment of samples with Ad-ATP12A is predicted to activate the TGF-β1 signaling pathway and subsequent downstream molecules.

Moreover, we used the IPA literature database to generate a heatmap of genes associated with the IPF signaling pathway (**Figure 3E**). A subset of this database was compared to the transcripts per million (TPM) data from the RNA-sequencing results for each of the mouse lung samples. We included ATP12A and MUC5AC in the list due to their relevance to this study. ATP12A expression was only present in lung samples of Ad-ATP12A expressing mice. This is consistent with the literature-supported observation that murine airway cells do not naturally express ATP12A. Lungs treated with BLEO showed increased levels of COL1A1 and COL3A1; a similar trend is observed for the matrix metalloproteins MMP2 and MMP7, with both being more prevalent in samples with induced ATP12A expression (**Figure 3E**).

### Potassium competitive proton pump blocker vonoprazan decreased ASL pH and TGF-β1 activation in IPF small airway epithelial cells

To investigate the potential roles of ATP12A in IPF small airways, we developed methods to culture human large and small airway epithelia at the air-liquid interface (ALI) by adapting a previously published method for culturing pig small airway cells (38). We examined if human small airway cells can form well-differentiated epithelium and express small airway-specific genes such as SCGB3A2, a highly expressed secretory protein in human small airways (39). In contrast to the minimal expression of SCGB3A2 in the large airway, there is an increased level of SCGB3A2 expression in the human small airway (**Figure 4B**). Both large and small airway epithelia have abundant acetylated α-tubulin positive ciliated cells (**Figure 4A and B**), indicating well-differentiated cells. Thus, we validated a human large and small airway culture model which preserves the native tissue properties. To test whether small airways from IPF patients maintain ATP12A expression in primary culture, we examined ATP12A expression by immunoblotting. As expected, robust ATP12A expression was detected in large airway epithelia but was not detected in normal small airway culture. Consistent with immunostaining of ATP12A in IPF lung tissue (**Figure 2**), there is increased ATP12A expression in IPF small airway culture (**Figure 4C and D**). As ATP12A is upregulated in IPF small airways, we tested the effects of proton pump inhibitors on ASL pH in IPF small airways. We recently reported that V-type ATPase containing ATP6V0D2 also can be contributing to the control of ASL pH in pig small airways (33). Interestingly, we found that V-type ATPase does not regulate ASL pH in human small airways from IPF lungs The V-type ATPase inhibitor Bafilomycin (BAF) does not affect ASL pH, but the ATP12A inhibitor ouabain inhibited ASL pH from IPF small airways (**Figure 4E**). In addition, VON by competitively inhibiting the binding of potassium ions to H^+^, K^+^-ATPases (including ATP12A) increased ASL pH in IPF small airway epithelial cells (**Figure 4E)**.

The overexpression of ATP12A enhanced BLEO-induced lung fibrosis and activation of the TGF-β1 signaling pathway (**Figure 3).** We hypothesized that ATP12A may play a role in latent TGF-β1 activation. To investigate the potential role of ATP12A in IPF small airways, we developed an assay to compare the activation of latent TGF-β1 in small airway epithelia from IPF and normal lungs. 100ng latent TGF-β1 (catalog 299-LT-005/CF, R&D) was applied to the apical surface for 24 hours; the activated TGF-β1 was quantified by a TGF-β1 ELISA kit. Small airway epithelial cells from IPF lungs activated almost 3-fold more latent TGF-β1 compared to small airways from normal lungs (**Figure 4F**). In addition, the pharmacological inhibitor VON reduced the rate of TGF-β1 activation by 25% (**Figure 4G**). Together, these data suggest VON may have anti-fibrotic roles *in vivo*.

### Inhibition of ATP12A by potassium competitive proton pump blocker, vonoprazan, reduced bleomycin-induced pulmonary fibrosis in mice expressing ATP12A

To confirm if ATP12A expression has a role in the enhancement of BLEO-induced pulmonary fibrosis in mice, we inhibited the action of ATP12A by using VON as a general gastric proton pump inhibitor. It has been reported that VON has a more potent and longer-lasting inhibitory effect on gastric acid secretion in animal models (40), compared to other traditional Proton Pump Inhibitors (PPIs) such as esomeprazole. VON is a K^+^-competitive acid blocker and binds to an extracytosolic domain of ATP12A or a gastric proton pump to block its function. Mice were administered VON daily via the oropharyngeal aspiration route for the duration of the experiment. Our data shows that VON significantly inhibited the effects of ATP12A expression on BLEO-induced pulmonary fibrosis. The lungs of these VON treated mice showed significant reductions in pulmonary collagen deposition (**Figure 5A, B, and C; Supplementary Figure 5A and B**), apoptosis in the alveolar epithelium (**Figure 5D and E; Supplementary Figure 5C**), airway mucous thickness (**Figure 5F and G; Supplementary Figure 5D**), and honeycomb cyst formation (**Figure 5H and I; Supplementary Figure 5E**) compared to the saline-treated group.

## Discussion

In the current study, we first investigated ATP12A expression levels in IPF, COPD, and normal human airways. Interestingly, we found clear evidence of ATP12A overexpression in IPF small airway surface epithelium and honeycomb cysts lining the epithelium. Moreover, MUC5B overexpression was observed in IPF small airways in association with ATP12A upregulation.

Overexpression of ATP12A in mouse lungs aggravates pulmonary fibrosis (PF) in a BLEO-induced experimental model of PF. Furthermore, the potassium-competitive proton pump blocker, VON, decreased profibrotic gene expression in IPF lung explant tissue. This data supports the hypothesis that ATP12A plays an important role in the pathogenesis of lung fibrosis. Our overall conclusion is that ATP12A mediated acid secretion in small airways and distal lungs impairs mucociliary transport, increases pH-dependent TGF-β1 activation, and enhances mucus accumulation and lung fibrosis; these effects can be blocked by the potassium-competitive proton pump blocker, VON.

### Role of ATP12A in lung diseases

ATP12A serves the main proton pump by moving protons out of epithelial cells in exchange for K^+^ to acidify ASL in large airway epithelia (31, 41, 42) (43, 44). ATP12A is required for CF lung disease development as it is responsible for proton secretions and the subsequent ASL acidification in the airways of CF patients (31, 45). Mouse lungs lack ATP12A expression; therefore, CF mice do not demonstrate all the characteristics of CF lung disease. Overexpression of ATP12A in CF mouse lungs decreases ASL pH, increases mucus viscosity, and impairs bacterial eradication abilities (31). Human and pig large airways express ATP12A, and therefore blocking ATP12A activity in CF human and pig large airways can enhance host defense (31). Additionally, ATP12A was shown to be a regulator of ASL viscosity, and ATP12A upregulation was associated with an increased ASL viscosity which was restored through the inhibition of ATP12A by ouabain (46). ATP12A overexpression may increase the mucus viscosity by impacting mucin biochemistry and structure through alterations in the luminal composition or the pH of secretory vesicles (47, 48). Airway epithelial secretions, including airway mucus, are prominent H^+^/HCO_3_^−^ buffers, and therefore ATP12A overexpression-driven proton secretion into the airways may increase airway mucus oxidation (49). This in turn increases mucin polymer disulfide cross-links that increase the viscosity and stiffen airway mucus (31, 45, 46, 50).

The hypersecretion and accumulation of mucus and mucociliary clearance impairment are common characteristics found in human IPF patients (21, 51). The major potential consequence of mucus accumulation and ciliary impairment, (resulting from the increased mucus viscosity), is the retention of inhaled foreign substances (air pollutants, microorganisms, …. etc.), which initiates chronic inflammation in alveolar regions and reduces lung functions (52). Also, mucus accumulation may promote mucus aspiration into distal airways, impairing the gas exchange process (53). Aspirated mucus may cause alveolar injury either by disrupting the surfactant surface tension properties or through interference with the interaction between alveolar type II cells and the underlying matrix (54); these alveolar injuries, over time, could lead to progressive fibroproliferation, microscopic scarring, and eventually IPF development (55, 56). Our findings demonstrate that ATP12A expressing mice showed significant accumulation of mucin and airway mucus plugging after BLEO exposure, which was prevented by inhibition of ATP12A activity by VON. Additionally, the co-expression of both ATP12A and MUC5B in the distal airways and the observed airway mucus buildup and plugging in mice expressing ATP12A after being injured with BLEO support the critical role of ATP12A in the pathogenesis of pulmonary fibrosis.

Honeycomb cysts (HCs) are clusters of fibrotic airspaces characteristic of the UIP observed in IPF. They are lined with pseudostratified columnar ciliated epithelium over cytokeratin 5 (KRT5) expressing basal cells and filled with mucus. Interestingly, we observed a significant number of honeycomb cysts (HCs) in mice expressing ATP12A 14 days after exposure to a single dose of BLEO by intratracheal instillation, which is not observed in the conventional bleomycin model. HCs were induced in the chronic phase after a single dose of BLEO exposure (51, 57). Further, it has been reported that MUC5B overexpression can elicit HC formation in the BLEO model and that the severity of pulmonary fibrosis and HC formation was correlated with the degree of MUC5B expression (51). In the current study, when ATP12A overexpressing mouse lungs were challenged with BLEO, it resulted in MUC5B overexpression in airways which induced early HC development.

To further confirm if the enhancement of pulmonary fibrosis observed in mice expressing ATP12A after BLEO exposure was ATP12A related, we inhibited the action of ATP12A through oropharyngeal administration of VON. VON is a novel oral potassium competitive acid blocker that competitively blocks the potassium binding site of H^+^-K^+^ ATPases (58, 59). Due to its higher pKa value, VON has a more stable inhibitory effect on gastric acid secretions than conventional proton pump inhibitors (PPI) and is commonly used in the treatment of peptic ulcers, gastroesophageal reflux, and Helicobacter pylori (H. pylori) eradication (58, 60–64). PPIs have been commonly used in IPF clinical treatment guidelines in many countries (65). In this study, VON administration significantly inhibited the synergistic effect of ATP12A expression on BLEO-induced pulmonary fibrosis in mice. These findings support the important role of ATP12A in the development of pulmonary fibrosis.

### TGF-β1 activation and its role in lung fibrosis

TGF-β1 is the essential profibrotic cytokine, and its over-activation mediates the development of PF (66–68). Latent TGF-β1 is synthesized by various cell types in fibrotic lungs including epithelial cells and myofibroblasts. The latent precursor form can be activated by various mechanisms: integrins, proteases, reactive oxygen species (ROS), mechanical stress, and low pH (69). Activated TGF-β1 binds to TGF-β1 receptors (TR-I/II), initiates kinase activity, and induces Smad2/3 phosphorylation. Phosphorylated Smad2/3 binds with Smad4 and translocates to the nucleus to mediate target gene expression. TGF-β1 can also function through non-Smad-dependent pathways to induce profibrotic effects. TGF-β1 can induce fibroblast to myofibroblast trans-differentiation, induce profibrotic gene expression, epithelial apoptosis, and epithelial-mesenchymal transition (EMT) (70–72). Epithelial cells usually form cell to cell contacts to ensure tissue integrity, but under certain circumstances such as injury, they lose these contacts along with their epithelial markers, subsequently gaining a fibroblast-like phenotype by expressing mesenchymal genes (73, 74). This process, known as EMT, is well studied in cancer development and metastasis (75, 76). We previously demonstrated that primary alveolar epithelial cells (AECs) cultured on a fibronectin surface undergo TGF-β1-mediated EMT (71). Furthermore, this process occurs *in vivo* for both IPF and experimental models of lung fibrosis (71). Thus, activation of latent TGF-β1 is critical for all downstream profibrotic effects. Interestingly, it has been reported that low pH can increase activation of latent TGF-β1 (77–80). We demonstrated that small airways from IPF lungs express upregulated ATP12A and there is more latent TGF-β1 activation that can be partially blocked by VON. This evidence supports the hypothesis that a lower pH microenvironment in distal lungs contributes to lung fibrosis pathogenesis.

### Apoptosis of distal airway/alveolar epithelial cells is involved in pulmonary fibrogenesis

As apoptosis is a highly regulated physiological process of cell removal, it plays a fundamental role in the homeostatic control of the cell population (81). An increase in the incidence of apoptosis within a given cell population can result in considerable cell loss over time. Therefore, up-regulated apoptosis is likely to account for the excessive loss of AECs or the failure to re-epithelize that is characteristic of pulmonary fibrosis. Studies in our lab and others using BLEO-treated rat and mouse models strongly support the role of epithelial apoptosis as a profibrotic event in fibrogenesis. Results obtained from studies on human lungs are consistent with those on animal models. Fragmented DNA (the hallmark of apoptosis) in bronchiolar and AECs was found in lung biopsies from patients with IPF (82). Similarly, our lab demonstrated a co-localization of fragmented DNA in the alveolar epithelium and alpha-smooth muscle actin (α-SMA), a marker for myofibroblast in biopsies from patients with IPF (83, 84). Collagen deposition and epithelial apoptosis induced by BLEO in rats and mice were blocked by ZVADfmk, a broad-spectrum inhibitor of caspase, one of the key enzymes mediating apoptosis (85, 86). In the current study, we detected increased apoptosis in ATP12A overexpressing mice treated with BLEO which can be prevented by VON. We speculate that lower pH in distal lungs enhances apoptosis, which is also a contributor to the fibrogenic process in both human lungs and animal models (87–89).

In summary, this study is conceptually innovative as it provides mechanistic insights into the role of small airways in the pathogenesis of IPF, which has been rarely investigated and poorly understood despite clinical data suggestive of its importance in IPF pathogenesis. The study focuses on the role of ATP12A in distal small airways from a PF animal model and human specimens. The findings of this study demonstrate the important role of small airways and ASL pH in the development of IPF and provide support for a novel therapeutic avenue to target the progressive fibroproliferation of this disease.

## Methods

### Human lung explant samples

Lung explant samples were obtained from 21 patients, (13 IPF and 8 COPD), undergoing lung transplantation. Pathologic assessment confirmed findings of advanced UIP in the IPF subjects and emphysema in the COPD group. Control samples (N=4) were obtained from donor lungs that were not suitable for lung transplantation (**Supplementary Table 1**).

### Induction of ATP12A expression in mice airways

We induced ATP12A expression in male C57BL/6J mice (aged 22±9 weeks) airways through intratracheal instillation of adenovirus subtype 5 encoding mouse ATP12A (ADV-253250, Vector Biolabs) with a dose of 10^8^ PFU/mouse in 50 µl of 2% carboxymethylcellulose (CMC) solution (Catalog 419273, Sigma Aldrich). Intratracheal instillation was performed under 3% inhalational Isoflurane anesthesia (NDC 11695-6776-2, Henry Schein). Two weeks after instillation mice were sacrificed to confirm the ATP12A expression by RT-PCR. In situ hybridization was used to confirm mRNA expression and immunofluorescence staining was performed to detect ATP12A protein.

### Effect of induced ATP12A expression in mice airways on bleomycin-induced pulmonary fibrosis

To investigate the role of ATP12A in pulmonary fibrosis, we used a BLEO-induced lung fibrosis mouse model. Only male mice were used in the current experiment due to their established sensitivity to BLEO (90, 91). Mice were randomly assigned into 6 groups; group 1, untreated control (control); group 2, treated with a single dose of adenovirus expressing green fluorescence protein (Ad-GFP, ADV-1060, Vector Biolabs); group 3, treated with a single dose of adenovirus expressing ATP12A (Ad-ATP12A); group 4, treated with a single dose of BLEO (BLM); group 5, treated with single doses of Ad-GFP and BLEO (BLM + Ad-GFP); group 6, treated with single doses of Ad-ATP12A and BLEO (BLM+Ad-ATP12A). Both Ad-GFP and Ad- ATP12A were administered to mice at a dose of 10^8^ PFU/mouse in 50 µl of 2% carboxymethylcellulose solution by intratracheal instillation. BLEO was administered to mice at a dose of 2 U/Kg body weight in 50 µl of saline by intratracheal instillation. Mice were observed daily for 2 weeks and sacrificed by cervical dislocation on day 15 under deep isoflurane anesthesia.

### Effect of inhibition of ATP12A by potassium competitive proton pump inhibitor, vonoprazan, on bleomycin-induced pulmonary fibrosis in mice expressing ATP12A

Mice were anesthetized by Isofluorane. BLEO (2U/ kg body weight) in 50 µl of saline or 50 µl saline alone, was instilled into the lungs through a catheter. The 50 μl dose was instilled at end-expiration, and the liquid was followed immediately by 300 μl of air to ensure delivery to the distal airways. As the viral vector was diluted in 2% carboxymethylcellulose solution to ensure airway delivery and transduction, we use 2% carboxymethylcellulose solution as the vehicle control in some groups. VON (100µM in 50ul saline) was administered by intratracheal instillation when Bleo or saline will be delivered. For the rest of time course, VON was administrated by pharyngeal instillation daily for 13 days. Mice were randomly assigned into 7 groups; group 1, untreated control; group 2, treated with single doses of BLEO and 50 µl 2% carboxymethylcellulose solution (BLM+CMC); group 3, treated with single doses of BLEO, 50 µl 2% carboxymethylcellulose solution, and 10 µM VON (TAK-438, MedChemExpress) in 50 µl of saline daily (BLM+CMC+VON); group 4, treated with single doses of Ad-ATP12A and BLEO (BLM+Ad-ATP12A); group 5, treated with single doses of Ad-ATP12A, BLEO, and 10 µM VON in 50 µl of saline (BLM+Ad-ATP12A+VON); group 6, treated with single doses Ad-GFP and BLEO (BLM+Ad-GFP); group 7, treated with single doses of Ad-GFP, BLEO, and 100 µM VON in 50 µl of saline (BLM+Ad-GFP+VON). Both Ad-GFP and Ad-ATP12A were administered to mice at a dose of 10^8^ PFU/mouse in 50 ul of 2% CMC solution by intratracheal instillation. BLEO was administered to mice at a dose of 2 U/Kg body weight in 50 µl of saline by intratracheal instillation. Mice were observed daily and administered either 100 µM VON in 50 µl of saline or saline by oropharyngeal aspiration daily for 2 weeks and then sacrificed on day 14. Lung tissues were harvested for histopathology, western blotting, RT-PCR gene expression, and RNA sequencing.

### Masson’s Trichrome, in situ cell death detection, in situ hybridization, and immunofluorescence staining of lung tissue

Human and mouse lung samples were inflated with a 4% w/v paraformaldehyde (PFA) solution in PBS (14190250, Thermo Fisher) then placed in 4% PFA overnight for fixation; then transferred to 15% w/v sucrose solution (207263, Fisher Chemical) in PBS for 48 hours at 4°C with gentle rocking, followed by 30% w/v sucrose solution for 72 hours at 4°C with gentle rocking. Lungs were then embedded in optimal cutting temperature (OCT) embedding medium (4585, Fisher Scientific), frozen at −80 °C, and sectioned into 10 μm thickness sections on Superforest plus gold microscope slides (15-188-48, Fisher Scientific) using LEICA CM 3050S cryostat (LEICA Biosystems). Masson’s trichrome staining was conducted using a trichrome staining kit according to the manufacturer’s protocol (ab150686, Abcam). Whole Masson’s trichrome-stained lung sections were visualized using Nikon Eclipse Ni-U bright-field microscope (Nikon, Japan) and analyzed by NIH ImageJ software through color deconvolution in a blinded manner(92).

### Histochemistry and immunofluorescence

For immunofluorescence detection of ATP12A, MUC5B, MUC5AC, and Krt-5 proteins; sections were fixed with ice-cold methanol, air-dried for 2 hours, hydrated in PBS for 15 mins, blocked with 5% BSA and 1 % goat serum in PBS for 2 hours, and then incubated overnight at 4°C with primary antibodies diluted in PBS containing 2.5% BSA and 0.5 % goat serum. The following antibodies and dilutions were used: rabbit anti-ATP12A (NBP2-92160, Novus Biologicals) at 1:500; rabbit anti-MUC5B (HPA008246, Sigma Aldrich) at 1:250; mouse anti–MUC5AC (45 M1) Alexa Flour 488 (NBP2-32732AF488, Novus Biologicals) at 1:500; rabbit anti-ProSP-C (AB3786, MilliporeSigma). Affinity purified rabbit polyclonal antibodies against recombinant N-terminal fragment (amino acids 13-102) of rat Atp12a were used to detect ATP12A protein in mice lungs at a dilution of 1:250 (29, 30). Following incubation with primary antibodies, tissues were rinsed 3 times in TBS-T and incubated with a solution of secondary goat anti-rabbit Alexa Fluor 488, goat anti-mouse IgG1 Alexa Fluor H&L 647 diluted at 1:1000 in PBS containing 2.5% BSA and 0.5 % goat serum for 1 hour in the dark. After further 3 washes in TBS-T, slides were counterstained with DAPI (2031179, Invitrogen) to stain cell nuclei then mounted with Vectashield Hardset antifade mounting media (H-1400-10, Vector Laboratories). For Keratin 5 detection in mice lungs, a secondary HRP antibody (ARH1001EA, AKOYA Biosciences) followed by Opal 520 fluorophore (FP1487001KT, AKOYA Biosciences) for signal amplification. For co-detection of ATP12A and MUC5B proteins expression in both human and mice lungs samples, the Opal 4-color Manual ICH kit (NEL810001KT, AKOYA Biosciences) was used according to the manufacturer’s instructions.

Confocal microscopy was performed using a laser scanning confocal microscope Nikon C2 (Nikon Instruments Inc.). DAB-stained slides were counterstained with hematoxylin (GHS332, Sigma Aldrich) and permanently mounted. Samples were visualized using a Nikon Eclipse Ni-U (Nikon Japan). For ATP12A protein expression analysis in human lung explants, the intensity of staining was randomly quantified in more than 10 images/explant. For MUC5B and Krt-5 staining analysis, both right upper and lower lung lobe sections were examined and analyzed in a blinded manner. Image analysis was performed using NIS element analysis and ImageJ (NIH) software.

### In situ cell detection

In situ detection of cells undergoing programmed cell death was performed utilizing the TUNEL reaction using in situ cell death detection kit, Fluorescein (11684795910; Roche Diagnostics) according to the manufacturer’s instructions. Before performing the TUNEL reaction, frozen lung sections were hydrated in PBS for 15 minutes, then incubated in permeabilization solution (0.1% Triton X-100 in 0.1% Na-citrate) for 10 minutes at 4°C. After washing with PBS, the sections were incubated in a humid chamber with TUNEL reaction mixture, prepared immediately before use, for 30 minutes at 37 °C in the dark. They were rinsed three times in PBS and counterstained with DAPI (2031179, Invitrogen), and then mounted with Vectashield Hardset antifade mounting media. Positive control sections were permeabilized and treated with DNase I 500 U/ml in 50 mM Tris-HCl buffer, pH 7.5, 10 mM MgCl_2_, 1 mg/ml bovine serum albumin, for 10 minutes at room temperature, before the TUNEL reaction. Negative control sections, after permeabilization, were incubated in label solution (without terminal transferase). Observations and photography were performed with a laser scanning confocal microscope Nikon C2 (Nikon Instruments Inc.). The nuclei of apoptotic cells showed green fluorescence due to the incorporation of fluorescein dUTP into the DNA strand breakages. 5-10 images per animal were randomly taken by confocal microscope and the percentages of apoptotic cells were determined blindly. Image analysis was performed using NIS element analysis and ImageJ (NIH) software.

### In situ hybridization

RNAScope detection was used to perform in situ hybridization according to the manufacturer’s protocol (Advanced Cell Diagnostics). Briefly, formalin-fixed frozen human and mice lung sections of 10 µm thickness were rehydrated in PBS for 15 minutes. Following citrate buffer (Advanced Cell Diagnostics) antigen retrieval, slides were rinsed in deionized water and\ immediately treated with protease (Advanced Cell Diagnostics) at 40°C for 30 minutes in a HybEZ hybridization oven (Advanced Cell Diagnostics). Probes directed against human or mouse ATP12A mRNA and control probes were applied at 40°C in the following order: target probes, preamplifier, amplifier; and label probe for 15 minutes. After each hybridization step, slides were washed two times at room temperature. Chromogenic detection was performed followed by counterstaining with hematoxylin (GHS332, Sigma Aldrich). Staining was visualized using a Nikon Eclipse Ni-U bright-field microscope (Nikon, Japan).

### RNA isolation and sequencing

RNA was isolated from samples using a TRIzol/chloroform extraction protocol. Briefly, tissue was homogenized in TRIzol™ Reagent (15596026, Invitrogen) and treated with chloroform. The sample was thoroughly mixed and incubated on ice before being spun down for 15 minutes at 12,000 x g at 4°C. After centrifugation, the aqueous phase was added to an equal volume of 70% ethanol and used as the starting material for the RNeasy Mini Kit (74104, QIAGEN). RNA concentration and quality were assessed using a NanoDrop™ One^C^ Microvolume UV-Vis Spectrophotometer (840274200, Thermo Scientific). Samples that met the quality control standards were submitted for Illumina sequencing. Sequencing was carried out by the Genomics Core at the Van Andel Institute.

### Differential expression analysis

Filtering settings for the initial edgeR analysis were an adjusted p-value ≤ 0.05 and |log 2 Fold Change| ≥ 1. Differential expression analysis of mouse samples was performed using the diffExp function found in the OCTAD R package (93). For all subsequent analyses, the filtering threshold was set to a p-value ≤ 0.01 and |log 2 Fold Change| ≥ 1. A more restrictive p-value threshold was used to isolate the most significant expression differences. To minimize the batch effect, mouse samples were split into their respective groups and compared to one another for downstream analysis (**Figure 3A**). PCA plots were generated using the NetworkAnalyst 3.0 visual analytics platform (94). Canonical pathways, upstream regulators, and signaling pathway predictions were generated with the QIAGEN Ingenuity Pathway Analysis tool (95). Bar plots of the common IPA pathways and upstream regulators were generated from the data by GraphPad Prism version 9.3.1. Gene expression changes were visualized using the GenePattern Heatmap Viewer available through the GenePattern analytical platform (96).

### Human large and small airway cell culture

We developed methods to culture human large and small airway epithelia at the air-liquid interface (ALI) using the similar protocol for pig small airway culture as published before (38). Studies included human donor lungs from two sources: 1) Normal donor lungs that did not meet the criteria for transplantation. These specimens are obtained and distributed for research purposes by the Michigan Gift of Life and Lung Bioengineering Inc. 2) Diseased lungs from patients with idiopathic pulmonary fibrosis (IPF) who are undergoing a lung transplant. These specimens are obtained and distributed for research purposes by the Spectrum Health system Lung Transplant Program under an IRB-approved protocol. All samples are de-identified and are not associated with any personal health identifiers. Briefly, large airway tissue is bronchi containing submucosal glands and complete cartilage rings support. Small airway tissue is dissected from the distal lung within 2cm from the edge of the lung parenchyma. Airway epithelial cells will be expanded using established methods developed to conditionally re-program airway epithelial cells (38). Briefly, large and small airway cells will be separately cultured in PneumaCult™-Ex Plus Medium (05040, STEMCELL Technologies) and maintained at 37°C with 5% CO2. After one passage of amplification, expanded epithelial cells were separated from feeder cells and seeded on collagen-coated semipermeable membranes (Corning) at a density of 10^6^ cells/cm^2^ and cultured at ALI at 37°C in a 5% CO2 atmosphere as previously described for 2–3 weeks. Large and small airway epithelial cultures were maintained in Small Airway Growth Media (CC-4124, Lonza) supplemented with 5% FBS and 10 ng/ml KGF (keratinocyte growth factor). All experiments were performed approximately 2 weeks after seeding on matched large and small airway epithelia that had been isolated from the same donor.

### RT-PCR

SuperScript™ IV VILO™ Master Mix (11756050, Invitrogen) with ezDNase enzyme treatment was used to synthesize cDNA from the extracted RNA. All samples of cDNA were diluted to a concentration of 1 µg/µL for the RT-PCR protocol. Primers for each gene of interest were ordered from Sigma-Aldrich (**Supplementary Table 2**), and a separate master mix was prepared for each primer set using the TB Green® Premix Ex Taq II (TLi RNaseH Plus) reagent kit (RR820A, Takara Bio). The final master mix contained primers at a concentration of 0.4 µM and a 1X concentration of TB Green® Premix Ex Taq II (TLi RNaseH Plus). A volume of 18 µL of the master mix was added to each well of a 96-well PCR plate (AB-0600-L, Thermo Scientific) with 2 µL of the previously synthesized 1 µg/µL cDNA. The reaction was run using the CFX96™ Real-Time System (Bio-Rad) and the results were quantified using the manufacturer’s software.

### Protein extraction and western blotting

To extract protein, frozen tissue was first thawed on ice before adding RIPA lysis buffer (786-489, G-Biosciences), with protease inhibitors (A32955, Thermo Scientific), to the sample. The sample was homogenized and spun down at 4°C at maximum speed for ten minutes. Total protein concentration was measured by performing a BCA protein assay (23235, Thermo Scientific) on the supernatant. For western blotting, a standardized concentration of protein was added to the 4X Bolt™ LDS Sample Buffer (B0007, Thermo Scientific). Except for the detection of collagen, 10X Bolt™ Sample Reducing Agent (B0009, Thermo Scientific) was added to each sample at a final concentration of 1X. Samples were then boiled at 95°C for ten minutes and loaded onto a Bolt™ 4 to 12%, Bis-Tris, 1.0 mm, mini protein gel (NW04122BOX, Invitrogen). Proteins were transferred onto a PVDF membrane using the iBlot™ 2 Transfer System (IB24001, Invitrogen). Membranes were blocked with 3% milk/TBST (T8793, Sigma-Aldrich) before overnight incubation with primary antibodies.

### Active TGF-β1 assay

The concentration of active TGF-*β1* was measured using the Human /Mouse /Rat /Porcine /Canine TGF-beta 1 Quantikine ELISA kit (DB100B, R&D Systems). Briefly, a sample volume of 50 µL was added to the coated ELISA plate containing the appropriate sample buffer. The plate was allowed to incubate at room temperature with the appropriate reagents, with a washing step in between each incubation. After the stop solution had been added, the absorbance was measured at 450 nm and 570 nm. To measure total TGFB1, samples must first be activated with a 1N HCL solution and then neutralized with 1.2N NaOH/HEPES before proceeding with the assay.

### Statistical analysis

Immunofluorescence, in situ cell death detection, and Masson trichrome data were analyzed by one-way ANOVA followed by Tukey’s multiple comparison post hoc test. RT-PCR and the number of honeycomb cysts data were analyzed by paired student’s t-test using Graph Pad Prism 9.3.1.

### Study approval

All protocols were performed in compliance with all relevant ethical regulations approved by local Institutional Review Boards; written informed consent was obtained from all patients who participated in the current study (Spectrum health IRB #: 2017-198). Studies using animals complied with all relevant ethical regulations. All animal studies were reviewed and approved by the Michigan State University Animal Care and Use Committee (Animal protocol approval #PROTO201900242).

## ONLINE SUPPLEMENTAL MATERIAL

**Supplementary Table 1.**
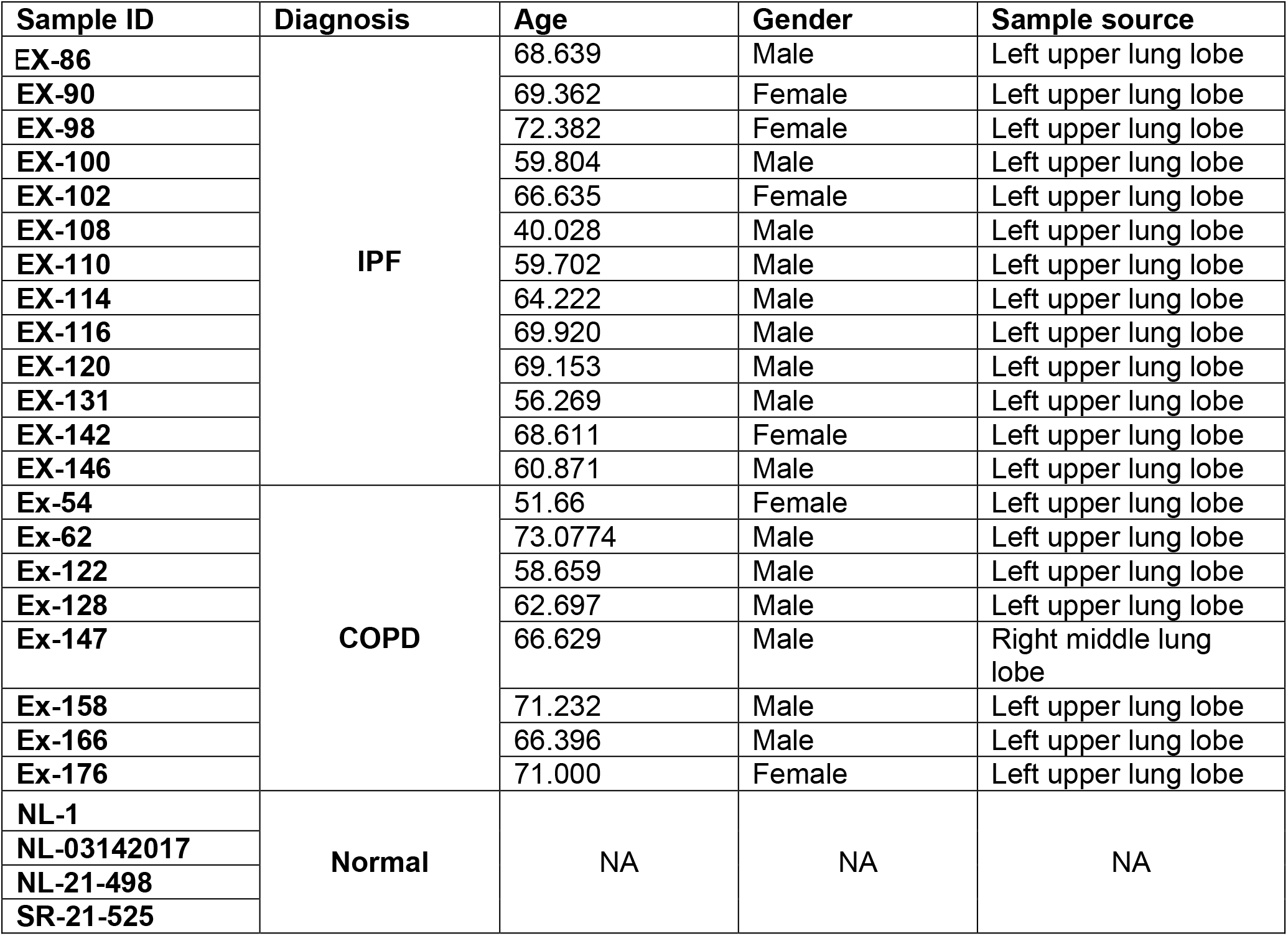
Summary of the clinical information of human samples used in the current study.

| Sample ID | Diagnosis | Age | Gender | Sample source |
| --- | --- | --- | --- | --- |
| EX-86 | IPF | 68.639 | Male | Left upper lung lobe |
| EX-90 |  | 69.362 | Female | Left upper lung lobe |
| EX-98 |  | 72.382 | Female | Left upper lung lobe |
| EX-100 |  | 59.804 | Male | Left upper lung lobe |
| EX-102 |  | 66.635 | Female | Left upper lung lobe |
| EX-108 |  | 40.028 | Male | Left upper lung lobe |
| EX-110 |  | 59.702 | Male | Left upper lung lobe |
| EX-114 |  | 64.222 | Male | Left upper lung lobe |
| EX-116 |  | 69.920 | Male | Left upper lung lobe |
| EX-120 |  | 69.153 | Male | Left upper lung lobe |
| EX-131 |  | 56.269 | Male | Left upper lung lobe |
| EX-142 |  | 68.611 | Female | Left upper lung lobe |
| EX-146 |  | 60.871 | Male | Left upper lung lobe |
| Ex-54 | COPD | 51.66 | Female | Left upper lung lobe |
| Ex-62 |  | 73.0774 | Male | Left upper lung lobe |
| Ex-122 |  | 58.659 | Male | Left upper lung lobe |
| Ex-128 |  | 62.697 | Male | Left upper lung lobe |
| Ex-147 |  | 66.629 | Male | Right middle lung lobe |
| Ex-158 |  | 71.232 | Male | Left upper lung lobe |
| Ex-166 |  | 66.396 | Male | Left upper lung lobe |
| Ex-176 |  | 71.000 | Female | Left upper lung lobe |
| NL-1 | Normal | NA | NA | NA |
| NL-03142017 |  |  |  |  |
| NL-21-498 |  |  |  |  |
| SR-21-525 |  |  |  |  |

**Supplementary Table 2.**
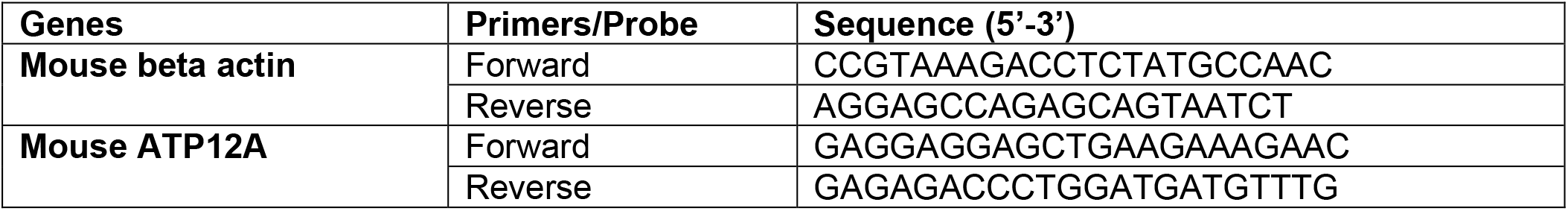
Forward and reverse primers sequances used for qRT-PCR.

**Supplementary Figure 1.**
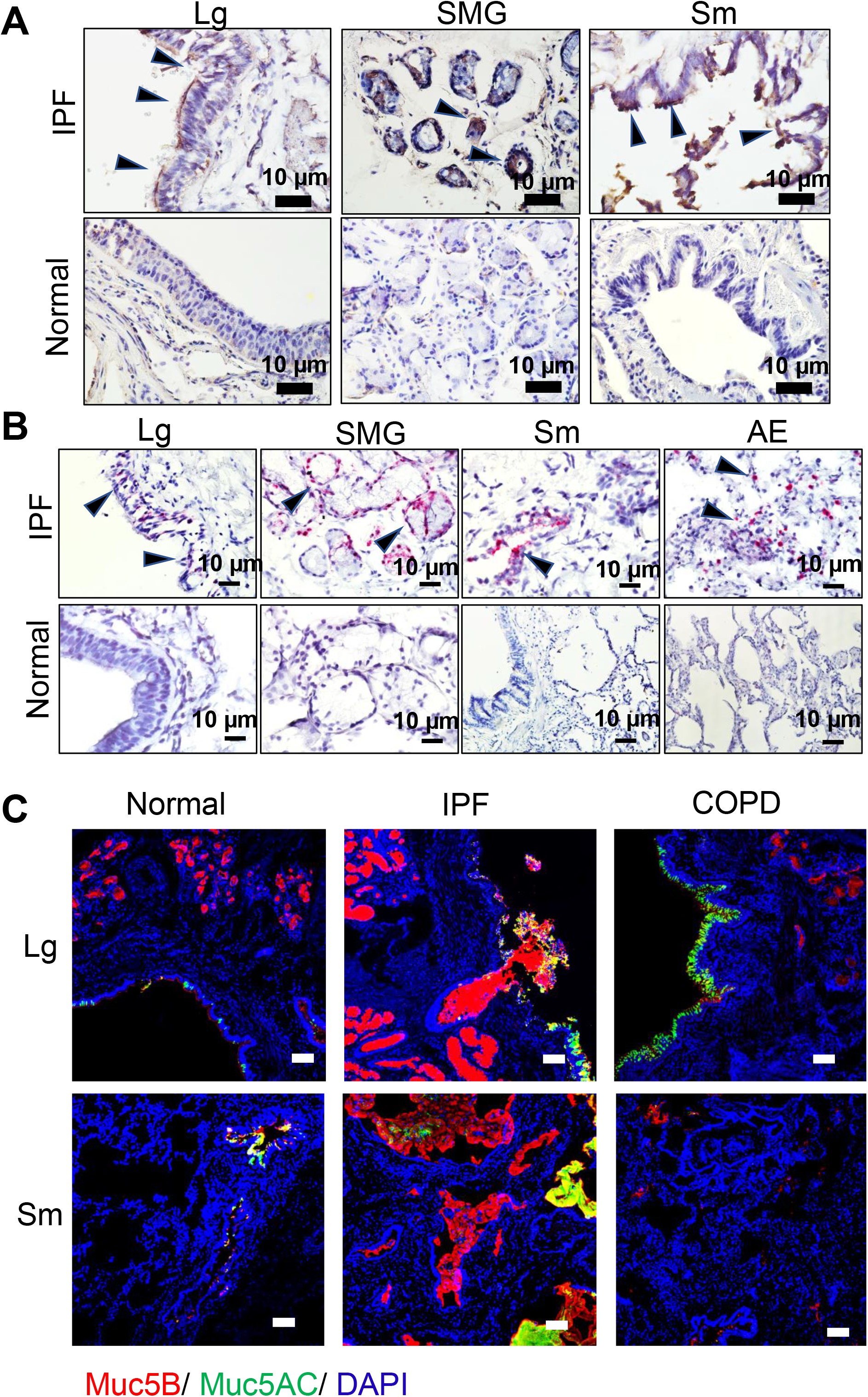
ATP12A, MUC5B, and MUC5AC proteins expression in human lung explants. **(A)** Representative brightfield microscope images showing immunodetection of ATP12A (brown, arrows heads) by immunohistochemistry. Nuclei were counterstained by hematoxylin (light blue). Scale bar: 10 µm. Images show large airways (Lg), submucosal glands (SMG), and small airways (SM) of normal (upper panel) and IPF (lower panel) human lungs. **(B)** Representative brightfield microscope images showing detection of ATP12A mRNA (pink, arrows heads) by in situ hybridization (ISH). Nuclei were counterstained by hematoxylin (light blue). Scale bar:10 µm. Images show large airways (Lg), submucosal glands (SMG), small airways (SM), the alveolar epithelium (AE) of normal (upper panel), and IPF (lower panel) human lungs. **(C)** Representative confocal microscope images showing immunodetection of MUC5B (red) and MUC5AC (green) by immunofluorescence. Nuclei were counterstained by DAPI (blue). Scale bar: 25 µm. Images show large airways (Lg) and small airways (SM) of normal, IPF, and COPD human lungs.

**Supplementary Figure 2.**
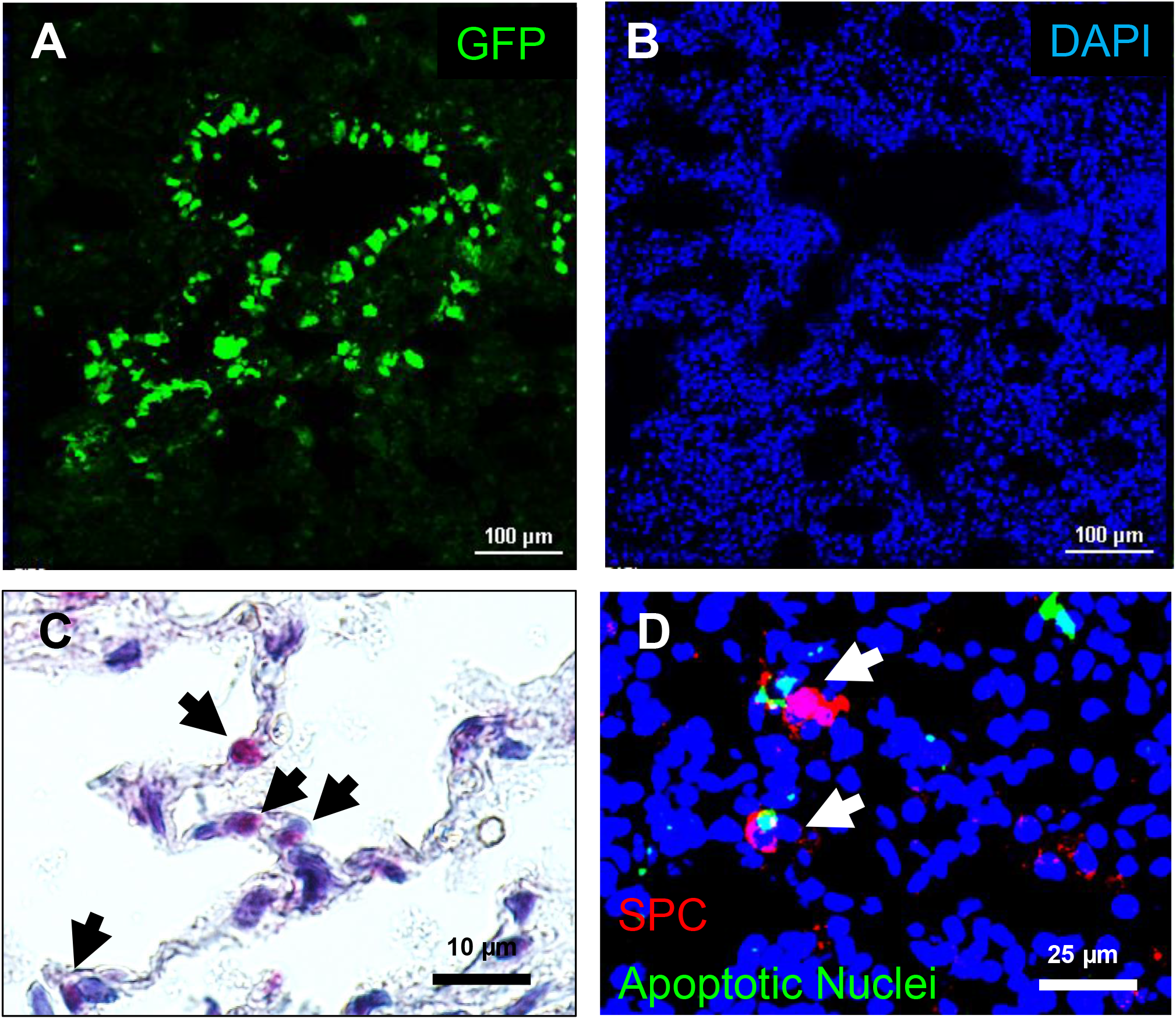
Detection of GFP expression and TUNEL^+^ cells in mice lungs. **(A and B) GFP was expressed in mouse lung treated with Ad-GFP. (A)** GFP protein (green color) was detected in mouse airways by confocal microscope imaging. **(B)** Nuclei were counterstained with DAPI in the same field as A. **TUNEL^+^ cells were detected in alveolar epithelial cells. (C)** Brightfield microscope image showing apoptosis in alveolar epithelial cells (pink, black arrows) of BLM+ATP12A group mice lungs. Nuclei were counterstained with hematoxylin. **(D)** Confocal microscope image showing apoptosis in type II alveolar epithelial cells (white arrows) of BLM+ATP12A group mice lungs. SPC (red, marker for type II alveolar epithelial cells) and apoptotic nuclei (green). Nuclei were counterstained with DAPI.

**Supplementary Figure 5.**
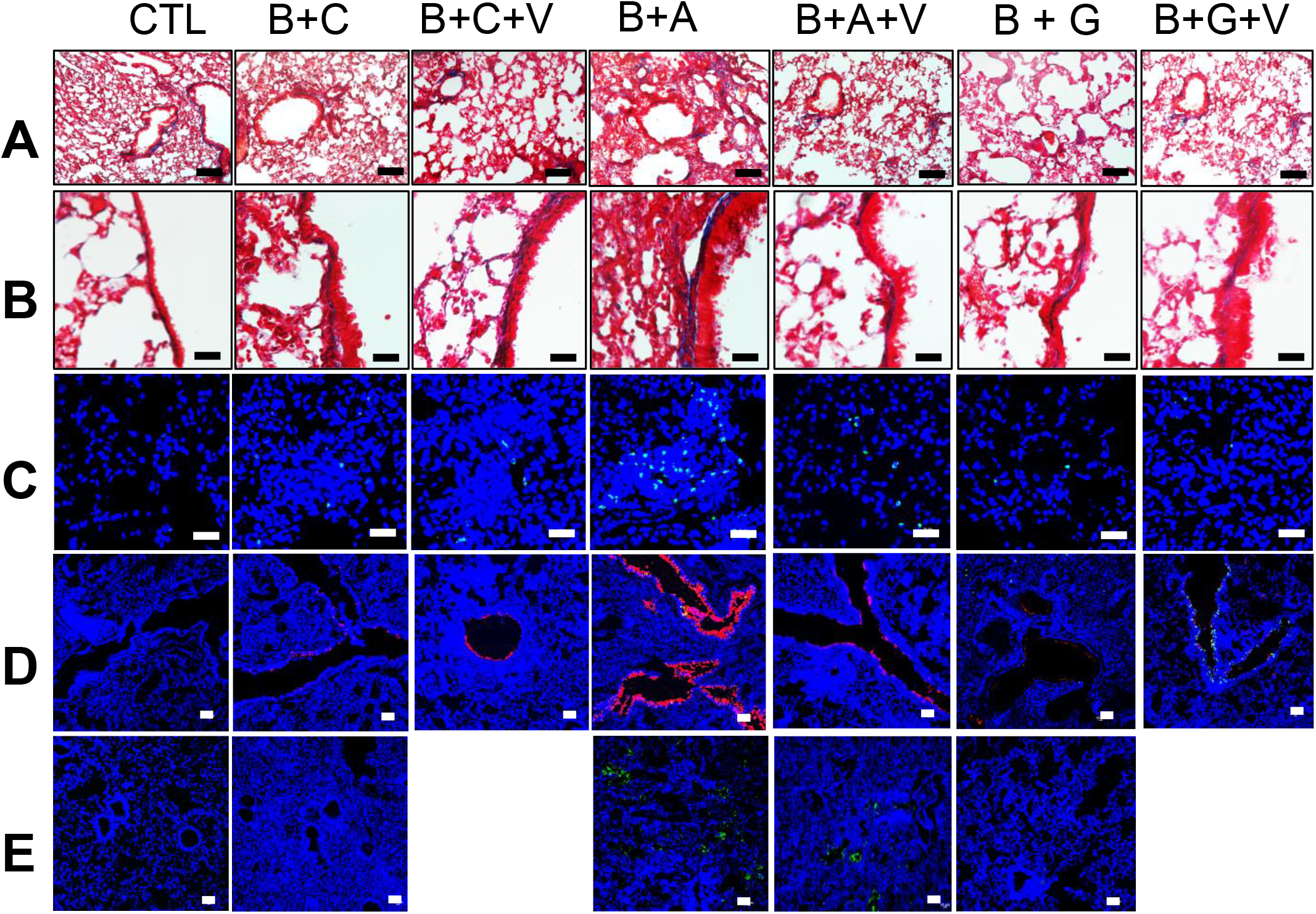
**(A)** Bright field microscope images of mice lung tissue stained with Masson’s trichrome showing collagen deposition (blue) in the lung, with the upper panel showing collagen deposition throughout the lung. Scale bar 100 µm; lower panel show collagen deposition in the parabronchial area. Scale bar 10 µm. **(B)** Confocal microscope images show cellular apoptosis of lung epithelial cells by TUNEL staining. Apoptotic cell nuclei are stained green. Scale bar: 25 µm. **(C)** Confocal microscope images show immunodetection of MUC5B (red) and MUC5AC (green) by immunofluorescence. Nuclei were counterstained by DAPI (blue). Scale bar: 25 µm.

